# Catabolism of lysosome-related organelles in color-changing spiders supports intracellular turnover of pigments

**DOI:** 10.1101/2021.02.22.432296

**Authors:** Florent Figon, Ilse Hurbain, Xavier Heiligenstein, Sylvain Trépout, Kadda Medjoubi, Andrea Somogyi, Cédric Delevoye, Graça Raposo, Jérôme Casas

## Abstract

Pigment organelles of vertebrates belong to the lysosome-related organelle (LRO) family, of which melanin-producing melanosomes are the prototypes. While their anabolism has been extensively unraveled through the study of melanosomes in skin melanocytes, their catabolism remains poorly known. Here, we tap into the unique ability of crab spiders to reversibly change body coloration to examine the catabolism of their pigment organelles. By combining ultrastructural and metal analyses on high-pressure frozen integuments, we first assess whether pigment organelles of crab spiders belong to the LRO family and, second, how their catabolism is intracellularly processed. Using scanning-transmission electron microscopy, electron tomography and nanoscale Synchrotron-based scanning X-ray fluorescence, we show that pigment organelles possess ultrastructural and chemical hallmarks of LROs, including intraluminal vesicles and metal deposits, similar to melanosomes. Monitoring ultrastructural changes during bleaching suggests that the catabolism of pigment organelles involves the degradation and removal of their intraluminal content, possibly through lysosomal mechanisms. In contrast to skin melanosomes, anabolism and catabolism of pigments proceed within the same cell without requiring either cell death or secretion/phagocytosis. Our work hence provides support for the hypothesis that the endolysosomal system is fully functionalized for within-cell turnover of pigments, leading to functional maintenance under adverse conditions and phenotypic plasticity. First formulated for eye melanosomes in the context of human vision, the hypothesis of intracellular turnover of pigments gets unprecedented strong support from pigment organelles of spiders.

## Introduction

How and why animals produce their colors are fundamental questions in biology. In cases where pigments are involved, they are usually synthesized and stored in specialized intracellular organelles (1). Bagnara and colleagues postulated in 1979 that all pigment organelles of vertebrates derive from a common primordial organelle (2). Since then, an increasing body of evidence has shown that pigment organelles, from mammal melanosomes to snake pterinosomes, belong to the lysosome-related organelle (LRO) family (3, 4). LROs intersect the endolysosomal system (5), which forms a complex and active network of membrane-bound compartments produced by endocytic and secretory pathways (6). Studying the subcellular aspect of LROs in relation to pigmentation is therefore a critical step to understand animal coloration.

Detailed investigations of intracellular processes and trafficking leading to the biogenesis of pigment LROs have been largely performed on mammalian melanosomes (3, 5). They revealed that skin melanocytes divert components of the endolysosomal system to progressively generate melanosome precursors derived from endosomes, bearing intraluminal vesicles and amyloid fibrils, and then mature pigmented melanosomes that are transferred to keratinocytes. Melanosome formation is controlled by a range of genes involved in the endolysosomal system of mammals (3, 5). The finding of homologous genes controlling pterinosomes, iridisomes and ommochromasomes of snakes and insects, among others, has led to a general model of pigment organelle formation (3, 4, 7). However, coloration does not only involve pigmentation phases but also bleaching phases that lead to pigment removal; a process that is far less understood, even for melanosomes (8).

While bleaching can result from death of pigment-containing cells, such as during peeling (9), this process is also compatible with pigment cells remaining alive (10). The latter phenomenon questions how pigment cells accommodate both the production and the removal of pigment organelles, as well as whether recycling pathways connecting the two phases exist. The occurrence of within-cell melanosomal degradation as a common physiological process required for melanin turnover has long been debated (8, 11–13). Here, turnover is defined as a pigmentation–depigmentation–repigmentation cycle at the cellular scale, without requiring the reuse of organellar materials. Evidence for physiological degradations of pigment LROs remains scarce [but see studies on differences in human skin pigmentation (14, 15)], which we attribute to the rarity of biological systems displaying active and concomitant production and removal of pigments within a single cell, and a lack of studies focusing on this phase.

Color-changing crab spiders can dynamically match the flower color on which they hunt. They do so by reversibly changing their body coloration between white and yellow via the metabolism of yellow pigments, thought to be ommochromes, in integument cells (16). This ability enables them to sit on flowers of various colors and wait cryptically for preys (17, 18). This non-model system is the best understood among the ones displaying morphological color changes in terms of pigment organelle metabolism, ranging from locusts to planarians (19, 20). During yellowing (*i.e.* anabolic phase), pigments are deposited within specialized intracellular organelles, whose intracellular origin remains undetermined (21). During bleaching, pigment organelles are thought to undergo an autocatalytic process and to recycle their membrane for another cycle of yellowing (10). However, ultrastructural evidence for this recycling process is scarce and we do not fully comprehend how both anabolism and catabolism of pigment organelles of crab spiders fit within intracellular trafficking pathways (7). Testing the hypothesis that these pigment organelles are members of the LRO family may help position them into the endolysosomal system, which in return might provide insights into the roles of the endolysosomal system in both pigmentation and bleaching phases, as well as in reversibility.

LROs are defined as being morphologically distinct from lysosomes, containing a subset of cell type specific contents necessary for their function and being in many instances secretory organelles. However, LROs also share features with lysosomes such as the presence of lysosomal associated membrane proteins, hydrolases, an acidic pH (at least temporary) and some can be accessible via the endocytic pathway (3, 5). In practice, the assignment of organelles to the LRO family is based on genetic defects associated to human diseases, such as Hermansky-Pudlak, Griscelli and Chediak-Higashi syndromes (3–5). However, such genetic dependence cannot be tested in spiders because they are in most cases not genetically tractable. Therefore, other markers of their secretory/endocytic nature should be assessed. LROs often bear ultrastructural signatures of their intracellular origin, such as endosomal/Golgi connections, intraluminal vesicles (ILVs), physiological amyloid fibrils and membrane tubulations (6). Conversely, LROs act as important regulators of metals by storing and releasing them in a dynamic manner (22). Specific techniques are required to retain such labile signatures as membrane tubulations and metals. For relatively thick tissues, this is best done by high-pressure freezing (HPF), which avoids artifacts produced by chemical fixatives (23). As an example, electron microscopy (EM) combined with HPF successfully revealed the morphological details of LROs like melanosomes (24) and Weibel-Palade bodies (25, 26) and their close contacts with other organelles (24). Given the sub-micron size of pigment organelles (21), detection of metals in their lumen can only be tackled by highly sensitive and spatially resolved nano-imaging methods (27), like scanning Synchrotron X-ray fluorescence (SXRF). SXRF is a state-of-the-art chemical imaging technique to map multiple trace elements with sub-part per million sensitivity down to a few tens of nanometers resolution (28–30). Therefore, to reveal the intracellular origin of pigment organelles and their catabolism, we exploited the combination of EM and SXRF on white, yellow and bleaching crab spiders fixed in a near-native state by HPF (see SI for the overall fixation quality; Fig. S1).

## Results

### Ultrastructure and Typology of Pigment organelles in Relation to Coloration

From an ultrastructural point of view, pigment organelles of white and yellow spiders mainly differed by the density and the organization of their intraluminal materials (Fig. S2). We classified pigment organelles preserved by HPF into six morphological types (Fig. 1A). The roundish a-type, corresponding to precursor organelles described in previous studies (10, 21), is present in both white and yellow spiders, although it was the main pigment organelle type in white ones (Table S1). HPF allowed to describe their internal content, consisting in a diffuse fibrillary material (Fig. 1A, a), with various stages of density (Fig. S2B). The roundish b-type, corresponding to mature pigment organelles also previously described (10, 21), is fully packed with electron-dense materials (Fig. 1A, b) and is predominant in yellow spiders (Table S1), reflecting its high pigment content. Contrary to a-and b-types, c- to f-types are only observed in yellow spiders (Table S1) and described for the first time. The c-type is an ovoid organelle characterized by several cores of variously dense materials forming pigment clusters of different natures (Fig. 1A, c, arrowheads; Table S1). The roundish d-type possesses a dense crown of material and an electron-lucent center (Fig. 1A, d; Table S1). The roundish e-type shows a reversed pattern with a core of dense material and a more diffuse material at the periphery (Fig. 1A, e; Table S1). Finally, the roundish f-type is characterized by a well-developed network of non-membranous fibrils resembling a ball of wool (Fig. 1A, f; Table S1). In addition, We observed that cores of c-types were similar to intraluminal contents of b-, d- and e-types (Fig. S3).

**Fig. 1.**
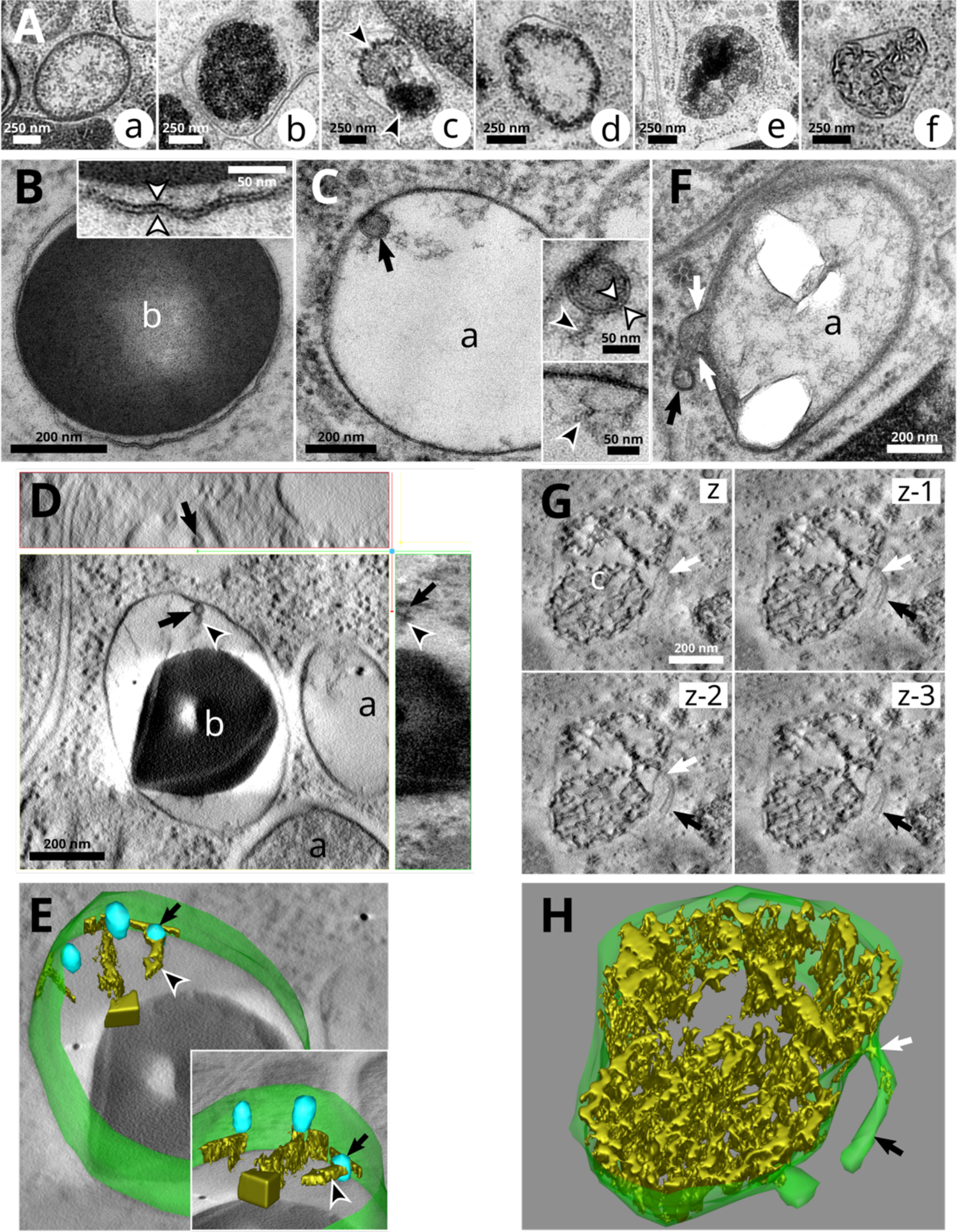
High-pressure freezing reveals fine ultrastructural features relating pigment organelles of crab spiders to lysosome-related organelles. (A) Pigment organelles are categorized into six types labelled a to f according to the morphology of their luminal content. (B) A b-type pigment organelle. Inset, double-leaflet structure (white arrowheads) of the limiting membrane. (C) An a-type pigment organelle containing an intraluminal vesicle (ILV, black arrow). Top inset, double-leaflet structure (white arrowheads) of the ILV membrane bearing fibrils (black arrowhead). Bottom inset, similar fibrils (black arrowhead) are attached to the limiting membrane. (D) Electron tomogram slices showing a fibril-bearing (black arrowhead) ILV (black arrow) attached to the limiting membrane of a maturing b-type pigment organelle. Top panel, XZ plane. Middle panel, XY plane. Right panel, YZ plane. (E) Three-dimensional reconstruction of the pigment organelle depicted in D showing its limiting membrane (green), ILVs (blue, black arrow) and fibrils (gold, black arrowhead). Inset, rotated view showing the sheet-like structure of fibrils. See Movie S1. (F) A membrane tubulation (black arrow) in continuity (white arrows) with the limiting membrane of an a-type pigment organelle. (G) Electron tomogram slices in the Z direction of a c-type pigment organelle bearing a membrane tubulation (black arrow) with a constricted neck (white arrow). (H) Three-dimensional reconstruction of the pigment organelle depicted in G showing the continuity between its limiting membrane and two tubules. See Movie S2. a, a-type pigment organelle; b, b-type; f, f-type.

### Fine 2D/3D Morphological Features of Pigment Organelles Revealed by HPF

The limiting membrane of pigmented organelles at various maturation stages show the typical 8-nm-thick bilayer structure (Fig. 1B, arrowheads in inset). All pigment organelle types could harbor intraluminal vesicles (Fig. 1C-E, arrows, and S4A-C) and tubulating membranes (Fig. 1F-H, arrows, and Fig. S4D). Membranes of intraluminal vesicles (ILVs) within pigment organelles were also well-preserved (Fig. 1C, white arrowheads in inset) and some were attached, or at least closely associated to the limiting membrane of pigment organelles (Fig. 1C and E, insets). One particular feature of these ILVs was the presence of a fibrillary material attached to their luminal face (Fig. 1C-D, arrowheads). Electron tomography (ET) revealed in three dimensions that those fibrils displayed a sheet-like structure expanding from the ILV surface into the pigment organelle lumen (Fig. 1E, arrowheads in the inset; Movie S1), similar to melanosomes (24). Tubules in continuity with pigment organelle membranes were also observed protruding into the cytoplasm (Fig. 1F-H, arrows). In Fig. 1F, the content within the tubule is denser than that of pigment organelles, indicating that it might carry substantial amounts of materials either from or to pigment organelles. In Fig. 1G-H, ET revealed a tubule (Movie S2) whose end was not connected to any other organelles (black arrow) and with a very narrow connection to the pigment organelle (white arrow).

### Analysis of Metals in Pigment Organelles by Correlative SXRF–STEM

We then investigated whether metal accumulated in pigment organelles using SXRF. We expected metals, if any, to be distributed locally and at low concentrations. To gain structural information during SXRF mapping, we took advantage of the presence of exogenously applied osmium (used as fixative during freeze substitution) to delineate structures at different scales, such as cuticle and cell layers (Fig. 2A), pigment cells (apposed to the cuticle; Fig. 2B) and organelles (Os-positive cytosolic structures; Fig. 2B-C, arrowheads). Native metal mapping reveals that those Os-positive structures differentially accumulate zinc (Zn), calcium (Ca) and, to some degree, cobalt (Co; Fig. 2C, insets and Fig. S6). The correlative SXRF-STEM approach confirms that Os-positive structures correspond to the dense intraluminal material of pigment organelles seen by EM (Fig. 2D, arrowheads). SXRF mapping at 50 nm/px even recapitulates the heterogeneous intraluminal organization between and within pigment organelles (Fig. 2D, arrowheads). Hence, mapping Os is an interesting alternative to visualize pigment organelles at various stages in case ultrastructural information is not readily available. Finally, by applying this correlative approach to native metals, we unambiguously showed that pigment organelles are the main cytosolic sites of metal deposition in pigment cells (Fig. 2E). A comparative analysis of white and yellow spiders by correlative SXRF–STEM further shows that metal accumulations in pigment organelle correlate with coloration (Fig. S6A-B). It further demonstrates that mature pigment organelles (b-types) are the preferred sites of Zn, Ca and Co depositions, while metal concentrations are markedly lower in a-, d-, e- and f-types (Fig. S6C-G; see Supplemental Results in SI for a detailed description).

**Fig. 2.**
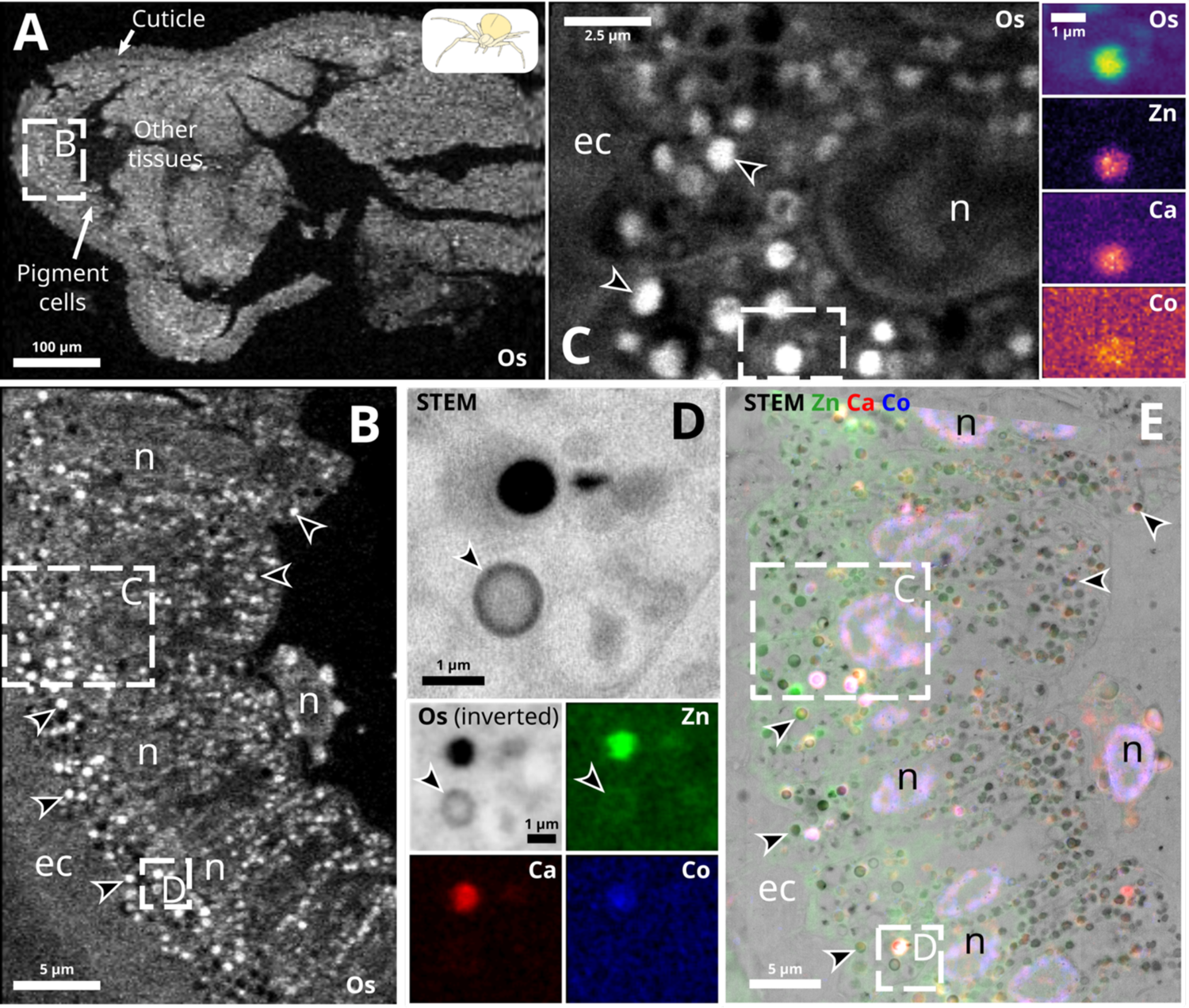
Correlative Synchrotron X-ray fluorescence and scanning transmission electron microscopy reveals metal accumulation in pigment organelles. (A-C) Hierarchical length-scale Synchrotron X-ray fluorescence (SXRF) of osmium (Os; white) enables the structural mapping of high-pressure frozen integuments, from tissues down to subcellular organelles. (A) Pixel size, 2 µm/px. Integration time, 10 ms. (B) Zoom on the region depicted in A. Pixel size, 500 nm/px. Integration time, 20 ms. (C) Zoom of the upper region depicted in B. Pixel size, 50 nm/px. Integration time, 20 ms. Inset, Os, zinc (Zn), calcium (Ca) and cobalt (Co) colocalize to discrete structures in the region depicted in C. Pixel size, 50 nm/px. Integration times, 20 ms (Os) and 300 ms (Zn, Ca and Co). (D) Scanning transmission electron microscopy (STEM) and SXRF (Os in black) of the lower region depicted in B show similar structures (e.g. black arrowhead). (E) Correlative SXRF-STEM of the region depicted in B. SXRF pixel size, 200 nm/px. Integration time, 200 ms. ec, endocuticle; n, nucleus.

### Intracellular Origin of Pigment Organelles

We next investigated the intracellular processes underlying pigment organelle biogenesis, maturation and degradation. Under the hypothesis that pigment organelles are LROs, both endosomal and secretory systems (*i.e.* Golgi and post-Golgi compartments) could contribute to their biogenesis (3). Therefore, we looked for ultrastructural evidence of either a Golgi or an endosomal origin of pigment organelles.

We spotted on several occasions elongated, swollen and stacked tubulo-saccular complexes that were in the close vicinity of a-type pigment organelles (Fig. 3A, asterisks). Those complexes were mainly found in the perinuclear region and sometimes in association with centrioles (Fig. S7). Fig. 3A-C show that these complexes are associated to endosomes (white arrow), microtubules (black arrowheads), as well as free tubules and coated and uncoated vesicles (white arrowheads), suggesting that they are regions of intense membrane trafficking. Strikingly, swollen cisternae of these complexes resembled the most electron-lucent a-types (Fig. 3A vs Fig. 3B and C). We next performed three-dimensional analyses by ET to further unravel the unique morphology of these complexes and their relationships with other compartments. Fig. 3D reveals that tubulo-saccular complexes are composed of distinct side-by-side compartments, each being a complex network of membranes forming tubules, saccules and elongated structures (Fig. 3E; Movie S3). Free vesicles are typically observed in between these compartments (Fig. 3D). A budding profile observed in Fig. 3 (arrowheads) indicates that tubulo-saccular complexes may be one source of free vesicles. The presence of an endosome with typical ILVs and an attached tubule (Fig. 3D, inset; white and black arrows, respectively; Movie S4) indicates that endosomal compartments could also generate some of the tubular and vesicular carriers observed in this region. Moreover, Fig. 3D shows that tubulo-saccular complexes are surrounded by a-type pigment organelles with different densities of intraluminal fibrils, which suggests that a-types mature in this part of the cell. Multiple vesicles can be seen in the vicinity of a-types, particularly at the interface with tubulo-saccular complexes (Fig. 3D). Strikingly, the swollen compartment from the tubulo-saccular complex harboring an attached vesicle contains elongated fibrils that emanate from the luminal face of the limiting membrane (Fig. 3F, black arrow), which is reminiscent of fibrils in a-types. The rather limited size of swollen compartments in tubulo-saccular complexes questions how a-types scales up to a micrometer and sometimes even more (see Fig. 3A).

**Fig. 3.**
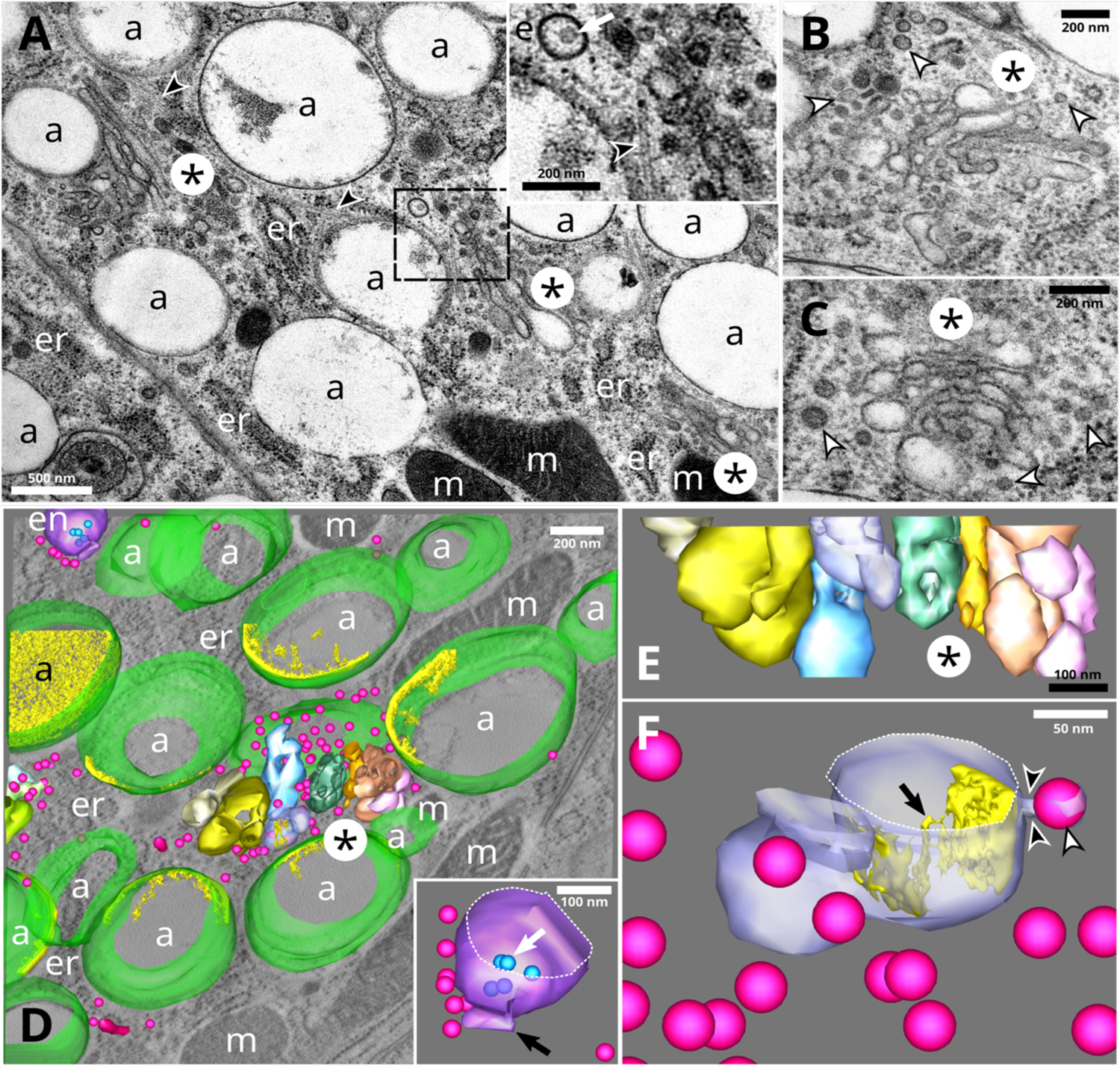
Maturing a-type pigment organelles are associated to tubulo-saccular complexes in regions enriched in endosomes, vesicles and tubular carriers. (A-E) Tubulo-saccular complexes (asterisks) are in the close vicinity of a-type pigment organelles. (A) Inset, zoom on an endosome (displaying an intraluminal vesicle; white arrow), vesicles and microtubules (black arrowhead). (B-C) Tubulo-saccular complexes showing swollen structures (asterisk), and numerous free coated and uncoated vesicles (white arrowheads). (D-E) Three-dimensional reconstruction of a tubulo-saccular region showing a-type pigment organelles (membrane in green and intraluminal fibrils in yellow), vesicles and tubules (fuschia), endosome (purple membrane and blue intraluminal vesicles) and tubulo-saccular complexes comprised of side-by-side compartments (different colors). Part of a second tubulo-saccular complex is seen on the left, surrounded by vesicles. Inset, close-up view of the endosome displaying intraluminal vesicles (blue, white arrow) and a tubule (black arrow). See Movie S3. (F) Details of one compartment from the main tubulo-vesicular complex in D showing the constricted neck (black arrowhead) of a budding vesicle (white arrowhead) and intraluminal materials (gold) forming fibril-like structures (black arrow) attached to the limiting membrane. See Movie S4. a, a-type pigment organelle; en, endosome; er, endoplasmic reticulum; m, mitochondria.

We therefore looked for ultrastructural markers of fusion events implicating pigment organelle precursors. We found many vesicles in the vicinity of tubulo-saccular complexes and a-types, although very few were directly attached to their membranes (Fig. 3). By contrast, we observed several homotypic fusion events between a-types, closely associated to microtubules (Fig. S8). However, we cannot exclude that those observations correspond to fission rather than fusion events. We reason that fission events of such big organelles would constrain their morphology, leading to their deformation near fission sites. Considering the relatively undisturbed aspect of limiting membranes, especially the absence of a constricted neck (Fig. S8A and C), fusion events seem more likely.

### Intracellular Degradation of Pigment Organelles

We were next interested in how c- to f-types were related to each other, hypothesizing that they represented catabolic stages of pigment organelle during bleaching rather than anabolic stages of the pigmentation process (*i.e.* intermediates between a-type precursors and mature b-types). To test this hypothesis, we captured yellow crab spiders, measured their body coloration and let them bleach overtime. Each spider was processed for HPF-FS and TEM analysis, representing a different state of bleaching (Fig. 4A). Under the hypothesis that c- to f-types are anabolic stages, we would expect them to occur mostly in yellow spiders and less in bleaching ones. Conversely, if they represent catabolic types, they should increase upon bleaching and vanish in fully white spiders (Fig. 4A). We found that c-, d-, f- and e-types (altogether referred to as intermediary types) were more abundant in bleaching spiders than in unbleached (*i.e.* yellow spiders, dominated by mature b-types) and fully bleached (*i.e.* white spiders, dominated by a-type precursors) spiders (Fig. 4A; see also Table S1). These results indicate that c-, d-, f- and e-types are transient organelles of the bleaching process rather than intermediary anabolic stages.

**Fig. 4.**
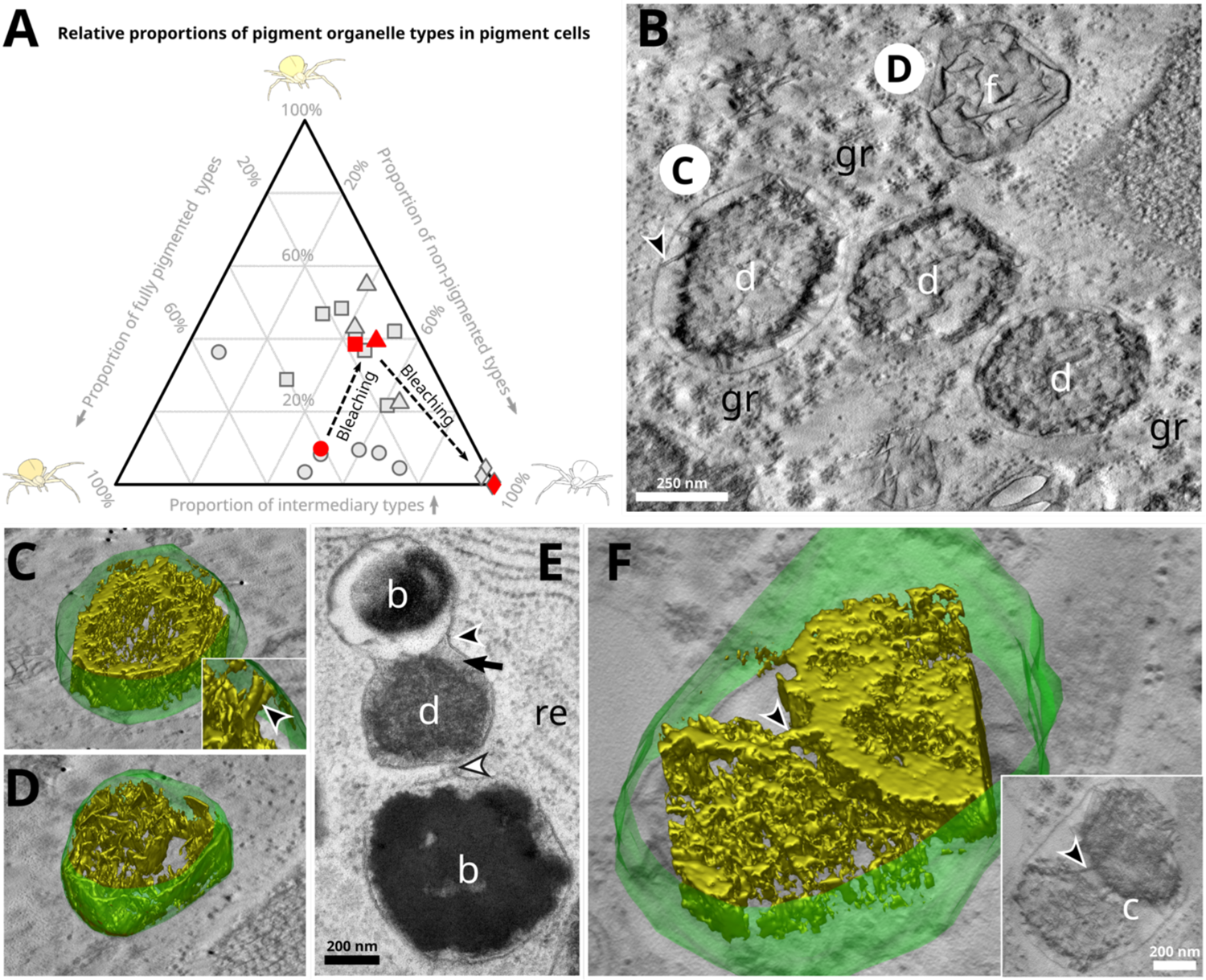
Degradation stages of pigment organelles during bleaching of crab spiders. (A) Ternary plot of the proportions in pigment organelle types for four bleaching spiders (circle, unbleached yellow spider; triangle, bleaching spider; square, more advanced bleaching spider; diamond, fully bleached white spider; see Materials and Methods). Between three to seven cells (grey symbols) were analyzed per individual. Geometric, rather than arithmetic, means (red symbols) were computed to preserve ratios between pigment organelle types. Intermediary types, c-, d-, e- and f-types; Non-pigmented types, a-types; Pigmented types, b-types. (B) Electron tomogram slice in the XY plane of d- and f-types in a bleaching spider. Black arrowhead, structure linking the internal content of a d-type to its limiting membrane. Gr, glycogen rosette. (C-D) Three-dimensional reconstructions of the limiting membrane (green) and the luminal content (isosurface rendering, gold) of two pigment organelles depicted in B. Inset of C, zoom showing a pillar-like structure (black arrowhead) linking the luminal content to the limiting membrane. See Movies S5 and S6. (E) Continuity (black arrowhead) and close-vicinity (white arrowhead) of limiting membranes suggest heterotypic fusion events between different pigment organelle types. (F) Three-dimensional reconstruction of a c-type cluster containing two pigment cores of different nature connected by a pillar-like structure (black arrowhead). See Movie S7. b, b-type pigment organelle; c, c-type; d, d-type; f, f-type.

We then investigated the ultrastructure of catabolic stages using 2D and 3D EM looking for markers of degradation. Fig. 4B-D show that intraluminal materials of d- and f-types are similar to each other, forming a core of dense material that extends in three dimensions and that connects the limiting membrane via pillar-like structures (Fig. 4C, arrowhead in inset; Movie S5). It suggests that f-types, whose fibrils are intertwined sheets (Movie S6), represent the latest stages of centrifugal degradation, in which only a backbone of sheets remains.

We finally examined the cluster-like c-type pigment organelles. We observe in several instances endosomes, multivesicular bodies and organelles displaying ultrastructural features of lysosomes in the vicinity of c- to f-types (Fig. S9). Fig. S3D shows a typical c-type cluster of five pigment organelles enclosed within a single limiting membrane. We observe in Fig. 4E a clear fusion event between two b- and d-types with continuous limiting membranes but with their respective lumens that have not mixed together yet (Fig. 4E, black arrowhead). Furthermore, a membrane extension seems to connect the middle e-type with another b-type (Fig. 4E, white arrowhead). Strikingly, the density and granular aspect of the e-type lumen resembles that of lysosome-like organelles previously described (compare Fig. 4E, black arrow and Fig. S9). The three-dimensional reconstruction of a c-type reveals that cores of materials are mostly independent of each other (*i.e.* there is no extent mixing of their contents; Fig. 4F), apart from some bridging regions with a pillar-like structure (Fig. 4F, arrowheads; Movie S7). They are reminiscent of the membrane-connecting structures observed in the inset of Fig. 4C, suggesting that these bridges are structural remnants of organelles before their incorporation into c-types.

## Discussion

### Pigment Organelles of Crab Spiders Are Lysosome-Related Organelles

Our morphological typology of pigment organelles optimally preserved by HPF matches with that of chemically fixed tissues (10, 21) but also allowed further insights into their fine morphological details. To the best of our knowledge, this is the first study employing correlative SXRF-STEM on such thick samples. This combination of nanoscale 2D/3D multimodal imaging techniques allowed us to observe features common to melanosomes, including intraluminal vesicles, sheet-like fibrils showing analogies with physiological amyloids (24), and zinc/calcium accumulation (30). Other studies described intraluminal vesicles and metals in pheomelanosomes (31, 32) and pterinosomes (33, 34), as well as various metals in insect ommochromasomes (35–37). These ultrastructural and chemical characteristics are shared by LROs independently of their nature and function (5, 22, 38), they hence associate pigment organelles of crab spiders to LROs.

### Pigment Organelles of Animals Likely Share an Endosomal Origin

We observed (unpigmented) a-type precursors of various sizes near tubulo-saccular complexes in the perinuclear region comprising endosomal compartments, microtubules, vesicles and tubules. Intraluminal fibrils similar to those of a-types were observed in a tubulo-saccular compartment. Altogether, these observations indicate that pigment organelles originate from these tubulo-saccular complexes (Fig. 5). It remains unclear whether tubulo-saccular organelles are modified Golgi apparatus (they share a stacked structure and numerous free vesicles) or endosomal compartments (they share tubular membranes and intraluminal fibrils). Both organelles are morphologically plastic (39, 40) and localize to the perinuclear region (6). Furthermore, both secretory and endocytic pathways, to which Golgi apparatus and endosomes belong respectively, are directly involved in the assembly of LROs, though at various steps of biogenenesis and maturation processes (3, 5, 41). We favor the hypothesis of an endosomal origin (Fig. 5) for three indirect reasons. First, pigment organelles possess ILVs, which have not been observed in LROs that originate from Golgi apparatus [e.g. Weibel–Palade bodies (25, 26)]. In line with this argument and to the best of our knowledge, inward budding has not been observed in Golgi compartments either. Second, the numerous striking similarities between pigment organelles and melanosomes, which derive from endosomes, imply that different origins for these two LROs are unlikely. Third, tubulo-vesicular complexes display morphological analogies with recycling endosomes, which are made of interconnected vesicles/tubules and contribute to melanosome biogenesis and maturation (42). Cytochemical experiments that could distinguish endosomal from Golgi compartments based on their enzymatic activities may clarify the intracellular origin of pigment organelles.

**Fig. 5.**
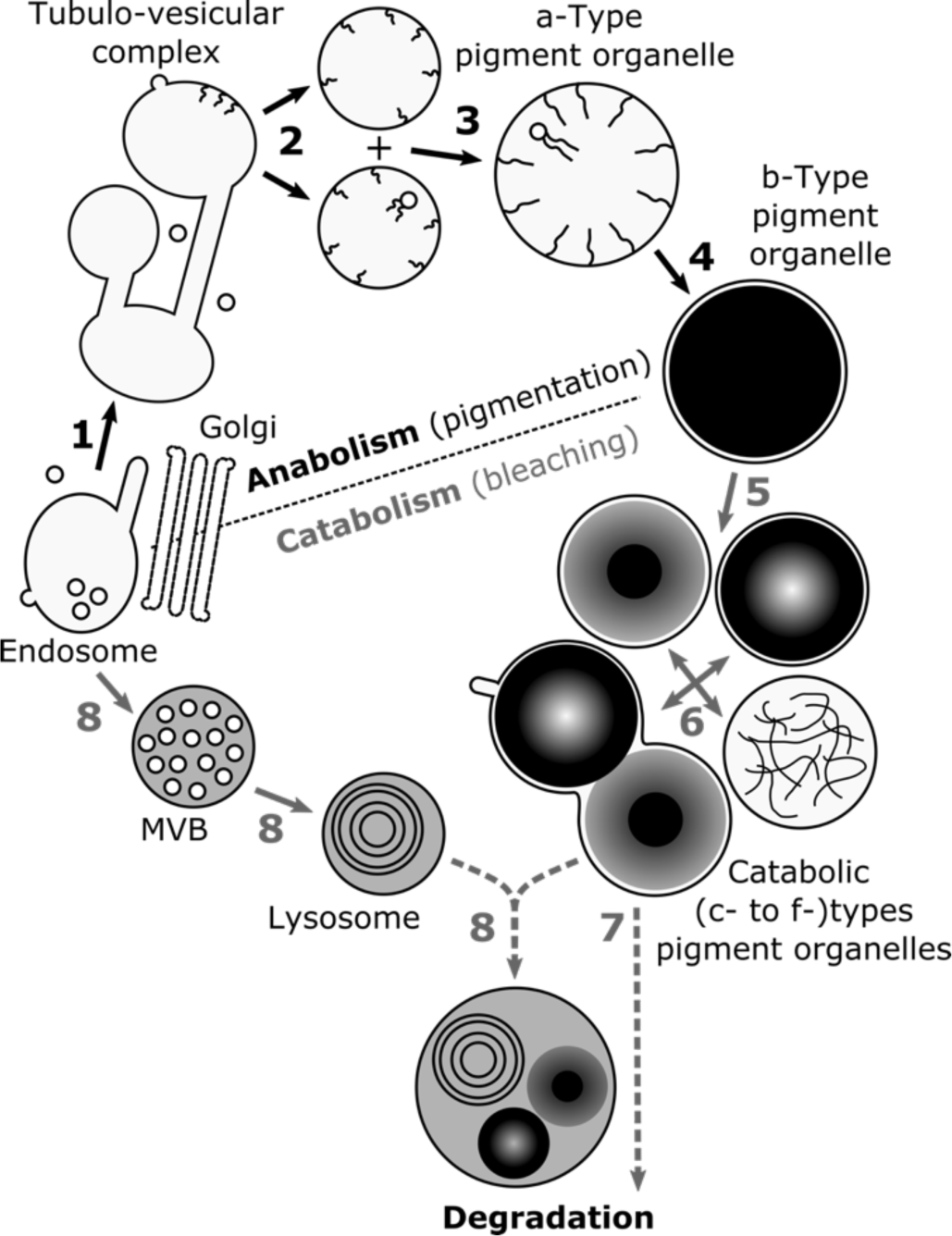
Working model of the endolysosomal-based pigment turnover leading to reversible color changes of crab spiders. (1) Tubulo-vesicular complexes originate from endosomes. (2) Tubulo-saccular complexes produce pigment organelle precursors (a-types) displaying intraluminal fibrils and vesicles. (3) Growth of a-types is sustained by homotypic fusion events. (4) Fibril densification and pigment deposition lead to fully pigmented and mature b-type organelles, thus to yellow coloration. (5) During bleaching, b-type content is variously degraded leading to catabolic d-, e- and f-types. (6) Catabolic types cluster together into c-types. Catabolic types are degraded by either (7) autocatalytic process or (8) fusion events with lytic compartments, such as lysosomes that also derive from endosomes through multivesicular bodies (MVB). The hypothetical nature of the last two steps is representede by dotted lines.

Overall, pigment organelles of crab spiders join the growing list of pigment LROs in distantly-related animals, including mammalian melanosomes (3), insect ommochromasomes and pterinosomes (43), fish pterinosomes (44), lepidopteran riboflavin and uric acid granules (45, 46), as well as snake melanosomes, pterinosomes and iridosomes (4). Our study therefore provides another piece of evidence that all pigment organelles, from invertebrates to vertebrates, share a common intracellular identity, as postulated 40 years ago by Bagnara and colleagues (2). In line with recent results (4), our work further suggests endosome is the primordial organelle of all pigment LROs (2).

### Pigment Organelle Precursors Likely Grow by Homotypic Fusion Events

Two mechanisms could be invoked regarding the growth of pigment organelle precursors (a-types): (i) numerous fusion events with vesicular and tubular carriers, and (ii) homotypic fusion events between early a-types. Although a large number of free vesicles and tubules were spotted in the vicinity of tubulo-saccular complexes and a-types, we did not observe many of these carriers attached to the limiting membrane of pigment organelle precursors. On contrary, homotypic fusion events, likely driven by microtubules, between a-types were more readily observed, indicating that these events are more frequent and thus predominant (Fig. 5).

### Intracellular Mechanisms of Bleaching in Crab Spiders in Relation to the Endolysosomal System

Based on the morphological similarities between catabolic types, their transient accumulation in bleaching spiders and the presence of the same metals but at lower concentrations, we suggest d- to f-types represent sequential degradative stages of mature b-types, culminating with their fusions into c-types (Fig. 5). This fusion process might be facilitated by the accumulation of Ca^2+^ in pigment LROs, since this ion is known to activate fusion of endolysosomal organelles (47).

How cells handle pigment LRO degradation intracellularly is a key question to understand bleaching of animals. Intracellular removal of pigments is classically mediated by three non-exclusive canonical mechanisms, namely secretion, autophagy and lysosomal degradation. Here, we focus on the last one (but see SI for further discussions of the two others).

For melanosomes, lytic activities were proposed to occur in two ways (8). First, lytic enzymes are delivered to melanosomes by their fusion with bona fide lysosome, the degradative organelle of the late endosomal pathway. Second, lytic enzymes that are intrinsically associated with melanosomes because of their lysosomal character get activated. In our analysis of bleaching crab spiders, we observed in several occasions catabolic pigment LROs with a dense granular lumen similar to multivesicular bodies and lysosomes, as well as catabolic pigment LROs fused with organelles morphologically similar to lysosomes. Lysosomes might hence potentially deliver their lytic content to pigment LROs (Fig. 5). The LRO identity of pigment organelles might facilitate their fusion with lysosomes by harboring endolysosomal fusion machineries (48), similarly to premelanosomes that exchange materials with lysosomes through kiss-and-run interaction thanks to PIKfyve kinase activity (49). Validation of lysosomal identity and fusion will require additional experimentation beyond the scope of this study. We also noted that catabolic d-, e- and f-types have nearly the same size as fully pigmented b-types and that catabolic types do not always show a lysosome-like dense lumen. Catabolic pigment LROs might hence be formed without extensive addition of lysosomal membrane and content, supporting the hypothesis that pigment LROs possess an autocatalytic activity (10). Such autocatalytic activity could stem from their lysosomal character, similar to other LROs like cytolytic granules in lymphocytes (50). Pigment LROs clusters (c-types) raise the interesting possibility that this autocatalytic activity might serve to degrade other pigment LROs by mimicking the effect of lysosomal fusion and thus delivering the lytic material of catabolically-active pigment organelles to unaltered ones. This contrasts with keratinocytes in white skin that accumulate melanosome clusters, morphologically similar to c-types but inactive, serving as pigment storage (51).

In addition to lytic pathways, recycling functions of the endolysosomal system should be considered. Endolysosomal organelles, including melanosomes, are known to recycle their content and membrane via tubulations (49, 52–54). We observed tubulations of a c-type, indicating that materials are actively recycled from catabolic stages during bleaching. Since metals are less concentrated in d- to f-types than in b-types, it is conceivable that tubules recycle metals for another turn of yellowing, bridging catabolism to anabolism of pigment LROs.

The presence of all pigment organelle types in yellow and bleaching spiders indicates that their anabolism and catabolism proceed simultaneously within the same cell, highlighting the great metabolic turnover of pigments within pigment cells (Fig. 5) (10). The numerous glycogen rosettes, mitochondria and lipid droplets observed in our electron micrographs and tomograms may be the fuel or the way to store energy from degraded pigments for another yellowing/bleaching cycle. Interestingly, glucose transporters have been shown to be involved in melanosome maturation (55), a process that might also be required for the yellowing of crab spiders after their bleaching. Further analysis on the link between glycogen content and coloration is warranted to test this hypothesis.

Overall, our results support the idea that pigment cells bleach via the *in situ* non-autophagic degradation of pigment LROs, which involve the continuous remodeling of both pigment organelle contents and membranes, presumably via lytic and recycling functions of the endolysosomal system.

### Within-Cell Turnover of Pigments: Physiological and Evolutionary Implication

We propose that, in the context of morphological color changes, the endolysosomal system is adapted and fully functionalized not only to produce and store pigments, but also to catabolize them without requiring cell death or phagocytosis (Fig. 5). Such a hypothesis of intracellular pigment turnover has been debated for melanosomes in mammal eyes (8, 11). It is still unclear whether melanosomes can be degraded intracellularly, even by other cells than melanocytes (8, 12, 51, 56, 57). Most degradative compartments containing melanin are autophagic and occur in pathological contexts, such as melanoma and vitiligo (8). Our results on pigment LROs of crab spiders imply that lysosomal degradation might exist for other pigments produced by the endolysosomal system. Such within-cell turnover of pigment organelles might allow organisms either to display phenotypic plasticity (e.g. slow reversible color changes; Fig. 5) or to cope with adverse conditions (e.g. photodegradation by free radicals or heavy metals) while maintaining cell functions. Because pigment LROs are ubiquitous in animals [this study and ref. (3, 4)], it further implies that the fundamental anabolic and catabolic functions of the endolysosomal system have been widely coopted during animal evolution to handle pigment-light interactions.

The fascinating color-changing ability of the non-model crab spider is thus in a unique position to provide key physiological understanding, otherwise very difficult to obtain, of LRO catabolism in the maintenance of light screening by human ocular melanosomes (8, 11).

## Materials and Methods

### Specimen Sampling

*Misumena vatia* (Clerck, 1757) crab spiders were collected in fields around the city of Tours, France, during summers 2018 and 2019. White and yellow individuals were sampled on plants harboring white and yellow flowers, respectively. They were kept individually in plastic vials no more than a week before dissection and fixation to avoid pigmentation loss.

To trigger bleaching of yellow spiders, individuals were left in plastic boxes covered with white cardboard and maintained in-door at room temperature. Spiders were fed once a week, at least two days before being further processed.

### Integument Fixation and Embedding

Crab spiders were sacrificed and fixed one at a time. For each individual, a few mm^2^ piece of integument was dissected from the opistosoma in Mg-containing Ringer’s solution supplemented with 2.2% glucose for isotonicity. High-pressure freezing (HPF) was performed with HPM100 (Leica Microsystems) or HPM Live µ (CryoCapCell) in 100% FBS serving as cryoprotectant. Freeze substitution was performed with AFS2 (Leica Microsystems) in anhydrous acetone containing 2% H_2_O / 1% OsO_4_. Samples were included in Durcupan AMC (EMS).

### Electron Tomography

Sections (300 nm) were cut with ultramicrotome UCT (Leica Microsystems). Random labeling was performed on both section sides with gold nanoparticles PAG 15 nm (AZU, Utrecht University, Netherlands). Single-tilt series with an angular range of −60° to +60° with 1° increment were imaged at 200 kV using transmission electron microscope Tecnai G2 (Thermo Fischer Scientific) equipped with TemCam-F416 4k CMOS camera (TVIPS) controlled by EM-Tools software (TVIPS). Alignment of projection images and tomogram computing (reconstruction-weighted back projection) were performed with Etomo in IMOD software (58). Modelisation was performed by manual contouring with 3dmod in IMOD software (59).

### Synchrotron X-Ray Fluorescence

Sections of 500 nm thickness were cut with microtome RM2265 (Leica Microsystems). They were deposited at the surface of distilled water drops lying on silicon nitride membranes (membrane size: 1 mm x 1 mm, membrane thickness: 1 µm, Silson). Membranes were quickly dried for a few seconds at 60 °C on a hot plate. Frames were glued on plastic washers to mount them on holders for Synchrotron X-ray fluorescence (SXRF).

Hierarchical length-scale SXRF imaging was performed at the Nanoscopium beamline of Synchrotron SOLEIL at Gif-sur-Yvette, France. Multi-elemental maps were acquired by fast continuous raster-scanning using the FLYSCAN scheme (60) with pixel sizes from 2 µm down to 50 nm and with exposure times from 300 ms to 10 ms per pixel. Full SXRF spectra were recorded for each pixel using two silicon drift detectors (KETEK) placed at ±120° to the beam direction (61). A 10.8 keV photon beam energy was used to excite all native elements from Al to Zn without exciting Os, which was added as a fixative during FS. The latter was specifically excited for morphological information by changing, on the fly, the beam energy to 11.4 keV.

Endogenous elements (10.8 keV) and osmium (11.4 keV) maps were aligned by using DaVis software (LaVision). The following parameters were used: “fill-up empty spaces,” “smoothing 2x at 3 px x 3 px” and “deformation with vector field.” Zinc signal was retained as marker for alignment because it presented similar features to osmium both in nuclei and cuticles, which allowed for a meaningful map correlation.

### Scanning Transmission Electron Microscopy

Frames analyzed by SXRF were removed from plastic washers and were introduced into a custom sample holder crafted to accommodate large silicon nitride frames. Osmium, without further sample staining, provided enough electron contrast to allow a direct comparison between STEM and SXRF maps. Scanning transmission electron microscopy was performed on a 200 kV field emission gun JEM-2200FS (JEOL, Tokyo, Japan) equipped with a bright-field STEM detector. A 10 µm condenser aperture was used and the camera length was set to 60 cm. In these conditions, the beam convergence and collection semi-angles were 6 and 6.6 mrad, respectively. Images were collected at a magnification of 40kX or 60kX depending on the experiment (corresponding pixel size 1.04 and 0.69 nm) using Digiscan II controlled by Digital Micrograph (Gatan, Pleasanton, CA, USA). To observe large areas and allow correlative analysis with SXRF images, mosaic STEM images were automatically collected using a lab-made script developped in Digital Micrograph.

### Correlative Synchrotron X-Ray Fluorescence and Scanning Transmission Electron Microscopy

Images of the same integument regions were sequentially acquired by SXRF and then by STEM. Correlative imaging was performed using eC-CLEM plugin (62) in Icy software (63). STEM images and osmium (or zinc) maps were set as target and transformed images, respectively. Non-rigid registration was done using natural fiducials markers, such as cuticle edges, nuclei and sample defects, which were easily and unambiguously observed in osmium maps and STEM micrographs.

### Image Analysis and Quantification

For color-changing experiments, spiders at different states of bleaching were processed by HPF-FS and TEM. To quantify proportions of pigment organelle types, only spiders showing the best preservation and ultrastructures were retained. Three to seven cells per spider were selected based on their section profile displaying both basal and apical poles and the nucleus. In each cell, pigment organelles were counted and classified based on their ultrastructural morphologies.

## Acknowledgments

We thank Teresita Insausti, Carole Labrousse, Suzanne Rochefort and Maryse Romao for their help, as well as Michael Marks for his comments on an early version of this manuscript. We acknowledge the Multimodal Imaging Centre at Institut Curie Orsay, for providing access to the cryo-electron microscopy facility. The authors would like to acknowledge the Cell and Tissue Imaging Platform – PICT-IBiSA (member of France– Bioimaging – ANR-10-INBS-32904) of the UMR144 of Institut Curie for help with electron microscopy. We acknowledge SOLEIL for provision of synchrotron radiation facilities (projects 20180104 and 20190205) and we would like to thank Gil Baranton for assistance in using the “Nanoscopium” beamline.

The authors declare no competing interests.

## Author Contributions

F.F. and J.C conceptualized the question; F.F., K.M., A.S., C.D. and J.C. designed research; F.F., I.H., X.H., S.T., K.M. and A.S. performed research; G.R. contributed analytical tools; F.F., I.H., X.H., S.T., K.M., A.S., C.D., G.R. and J.C. analyzed data; F.F. and J.C. wrote the paper; All authors reviewed and edited the paper.

## Supplemental Information – Figon *et al*

### Supplemental Materials and Methods

#### Color Measurements

Body coloration of crab spiders was quantified by measuring reflectance spectra of their integuments as described in (1). For each spider, and time point during the course of color change, color indexes were averaged from three reflectance spectra taken successively. Body coloration was measured the day after capture and no more than a day before processing spiders for further imaging.

#### Reagents and Solutions

Glucose, Fetal Bovine Serum (FBS), magnesium chloride, sodium chloride and potassium chloride were purchased from Sigma-Aldrich. Uranyl acetate, lead citrate, Durcupan ACM, anhydrous acetone and osmium tetroxide were purchased from EMS.

#### Transmission Electron Microscopy

Ultra-thin sections (70 nm) were cut with ultramicrotome UCT (Leica Microsystems), stained with uranyl acetate and lead citrate and imaged with a Tecnai Spirit electron microscope (Thermo Fisher Scientific, Netherlands) equipped with à QUEMESA CCD camera (EMSIS) and iTEM software (EMSIS).

#### Image Analysis and Quantification

The different stages of pigment organelles observable in TEM images were manually contoured in several cells per individual. To calculate the proportions of pigment organelle types, three yellow and three white spiders captured the same day and cryofixed at the same time were used as biological replicates. Proportions were reported as the mean ± standard deviation of the three spiders per color. Morphometric data (maximum Feret diameter as a proxy of size and elongation ratio as a proxy of ellipticity) were automatically computed using Icy software and were reported as the mean ± standard deviation of 666 and 779 organelles observed in the three white and the tree yellow spiders, respectively.

For endogenous metal quantifications, backgrounds of SXRF maps were removed by subtracting to each pixel the average pixel value within a rectangular region comprising only resin using Fiji software (2). Structures positive for osmium were manually selected in the aligned Os maps using round ROIs of 800 nm diameter in Icy software. ROIs were copy-pasted onto endogenous metal maps. Mean pixel values within ROIs were calculated and extracted using Icy software.

#### Statistical Analysis

All statistical analyses were performed using R software (3). Principal component analyses and hierarchical clustering were done using FactoMineR library. Compositional analyses (ternary plot) of pigment organelle types in color-changing experiments were performed using RobCompositional library. Statistical differences between SXRF populations were tested first by non-parametric Kruskal-Wallis tests and then by pairwise Wilcoxon tests. Only p-values below 0.05 were considered as statistically significant.

### Supplemental Results

#### Metal Accumulation in Relation to Coloration

We investigated whether the morphological diversity of pigment organelles we described was associated to a variation in metals. First, we compared the distribution of metals in white and yellow integuments using Os as a structural marker of pigment organelles (Figure S6A and B). Both types of integuments showed high densities of pigment organelles (Os-positive structures) but yellow ones harbored higher amounts of Os (Figure S6B), in agreement with the presence of more pigmented organelles binding more Os in these tissues. While in white tissues, nuclei accumulate either similar or higher amounts of Zn, Ca and Co than those in yellow tissues (Figure S6A and B), pigment organelles of yellow tissues accumulate higher amounts of those metals (Figure S6B). Zn is seemingly present in some pigment organelles of white integuments, albeit in fewer amounts than in yellow tissues. These results suggest that metals are preferentially associated with yellow coloration.

We then investigated whether there was a relationship between pigmentation status at the level of pigment organelles and the presence of metals in yellow integuments. First, we performed correlative STEM-SXRF (Fig S6B-D) that qualitatively showed that more pigmented organelles (b- and c-types) accumulate more metals (Figure S6C-D). Next, we quantified the amount of metals within each Os-positive structure of Figure 4B and we performed a multivariate analysis by principal component analysis (Figure S6E). The first cluster of pigment organelles was characterized by low concentrations of metals, including Os (Figure S6F, cluster 1), indicating that the less pigmented organelles (based on Os signal) accumulated less metals. Second and third populations were mostly similar in terms of pigmentation but differed in their concentrations of native metals (Figure S6F, clusters 2 and 3). Zn, Co and Ca were 1.40, 1.67 and 1.80 times more concentrated, respectively, in population 3 than in population 2.

We then asked whether this heterogeneity in metal distribution was related to morphological diversity. A correlative SXRF-STEM experiment was performed to categorize each Os-positive structure within one of the pigment organelle types described in Figure 1E. Figure S6G validates that Os signal recapitulates the pattern of intraluminal density in pigment organelles as population 1 is enriched in a-, d- and e-types. Populations 2 and 3 do not segregate to distinct morphological types, they are rather enriched in b- and c-types, as well as in organelles that were intermediate between a- and b-types. Nonetheless, population 1 is clearly underrepresented in fully pigmented b-types (Figure S6G), which further supports a link between pigmentation status and metals.

### Supplemental Discussion

#### Subcellular Mechanisms of Bleaching in Crab Spiders

Recycling pigment organelles back by autophagy for another cycle of yellowing after the complete catabolism of their content has been proposed for crab spiders (4). This hypothesis is not supported by our observations because the electron-lucent vacuoles associated to ER whorls and previously hypothesized to be recycled pigment organelles (4) are actually lipid droplets. Regarding the pigment secretion hypothesis, we never observed pigment organelles in the extracellular space nor their fusion with the plasma membrane, precluding thus their exocytosis.

### Supplemental Figures

**Fig. S1.**
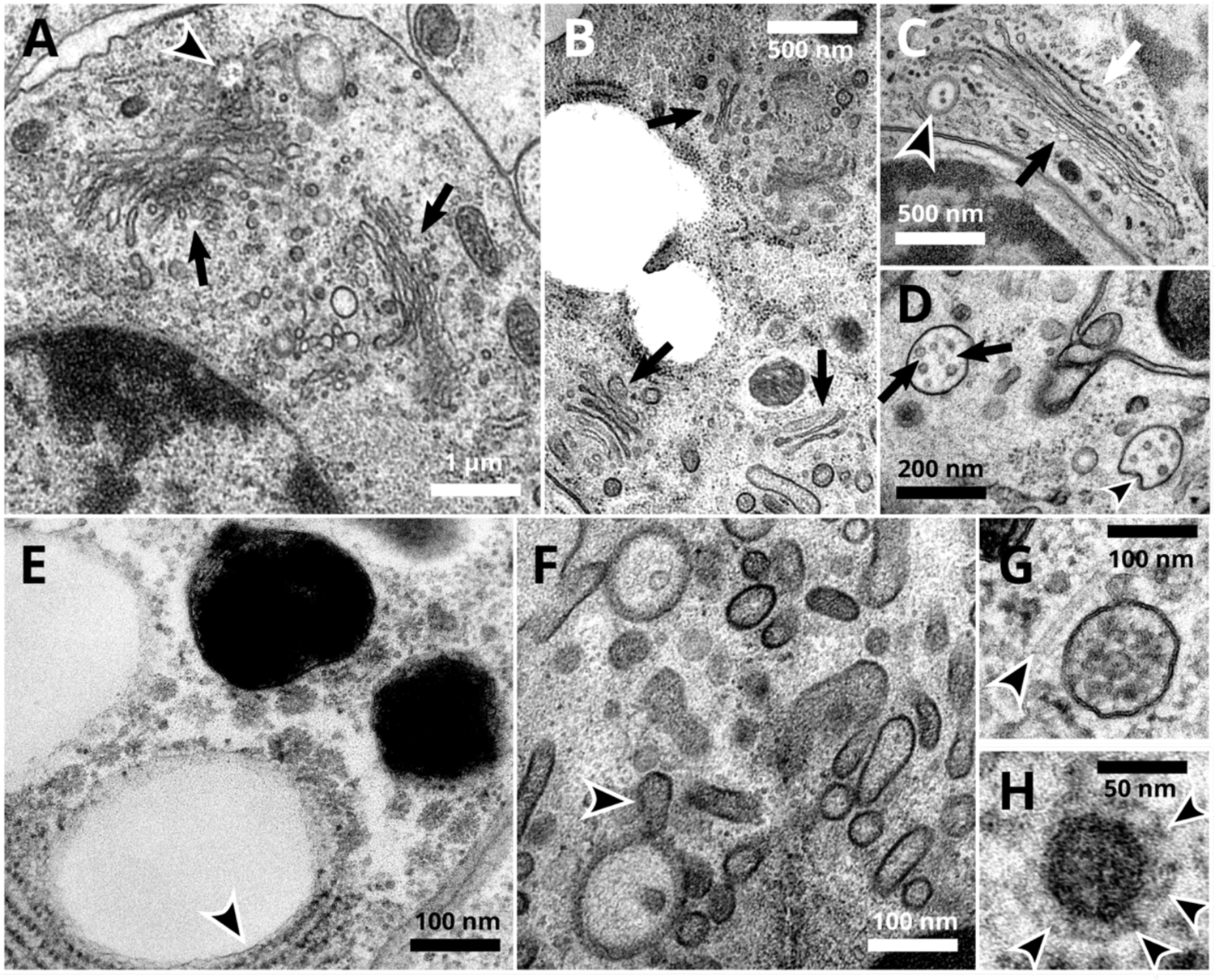
Ultrastructure of crab spider cells preserved by high-pressure freezing and freeze substitution. (A) A non-pigment cell showing well-preserved nucleus, mitochondria, Golgi apparatus with clear stacked structures and budding profiles (arrows) and a closely associated endosome (arrowhead). (B) Multiple Golgi apparatus (arrows) with neat stacked structures within a non-pigment cell. (C) A stacked and budding Golgi apparatus showing distinct cis-(white arrow) and trans-(black arrow) sides in a non-pigment cell. An endosome harboring intraluminal vesicles (arrowhead) is closely associated. (D) Two endosomal compartments with multiple intraluminal vesicles (ILVs) and fibrils (arrows). The limiting membrane of one of these endosomal compartments is budding inward (arrowhead), likely forming a new ILV. (E) A lipid droplet closely associated to endoplasmic reticulum (arrowhead). (F) Endosomes and tubules within a non-pigment cell. An endosome display a tubulating membrane (arrowhead). (G) A microtubule (arrowheads pointing to its ends) closely associated to cytosolic vesicles and to a budding multivesicular body. (H) A cytosolic vesicle displaying a coat of proteins (arrowhead) resembling clathrin.

**Fig. S2.**
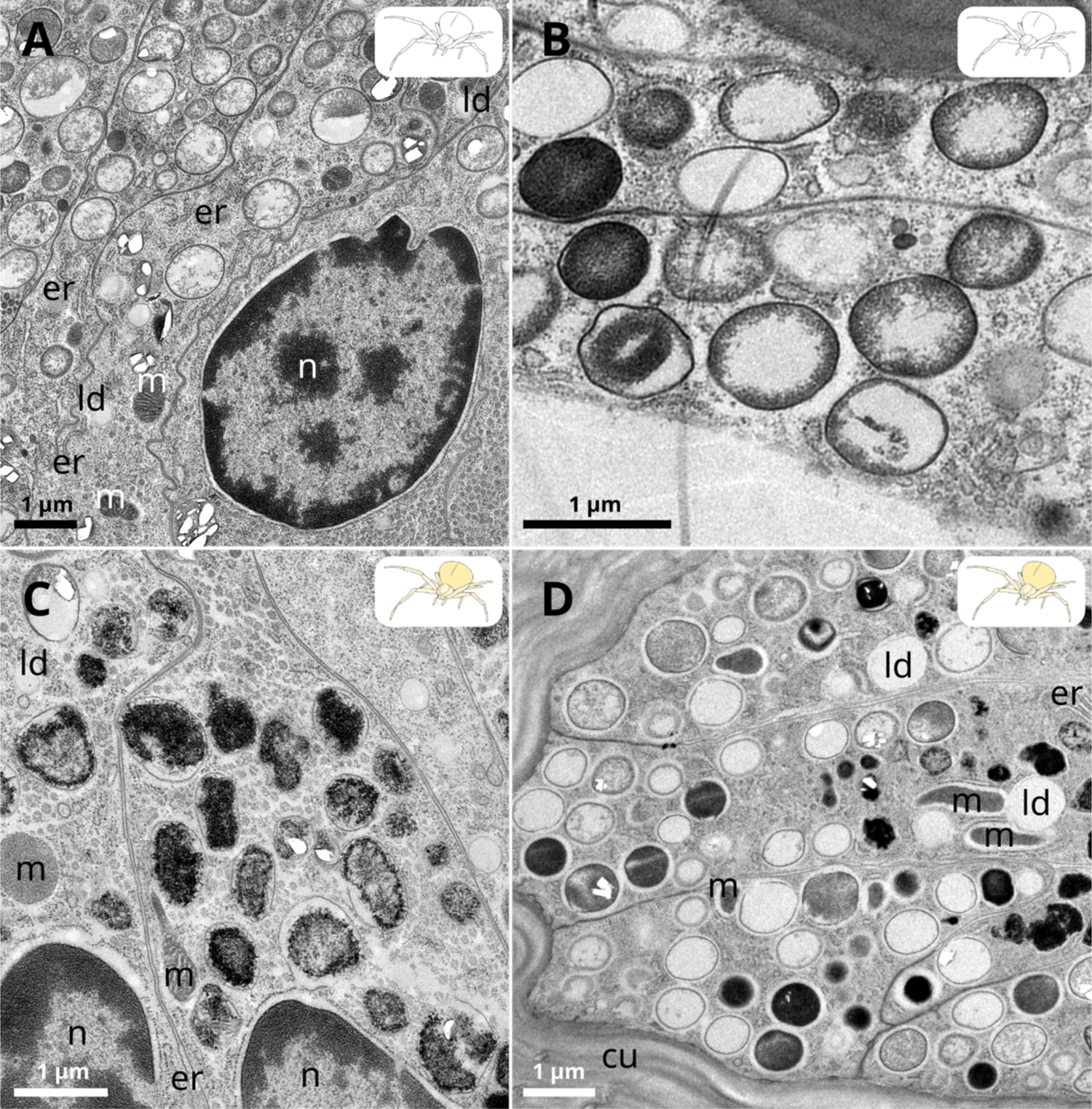
Ultrastructural diversity of pigment organelles. (A) Early stages of pigment organelles in a white crab spider. (B) Pigment organelles at different stages of filling in a white spider. (C) Pigment organelles observed only in a yellow crab spider. (D) Variety of pigment organelles observed in yellow spiders. cu, cuticle; er, endoplasmic reticulum; ld, lipid droplet; m, mitochondria; n, nucleus.

**Fig. S3.**
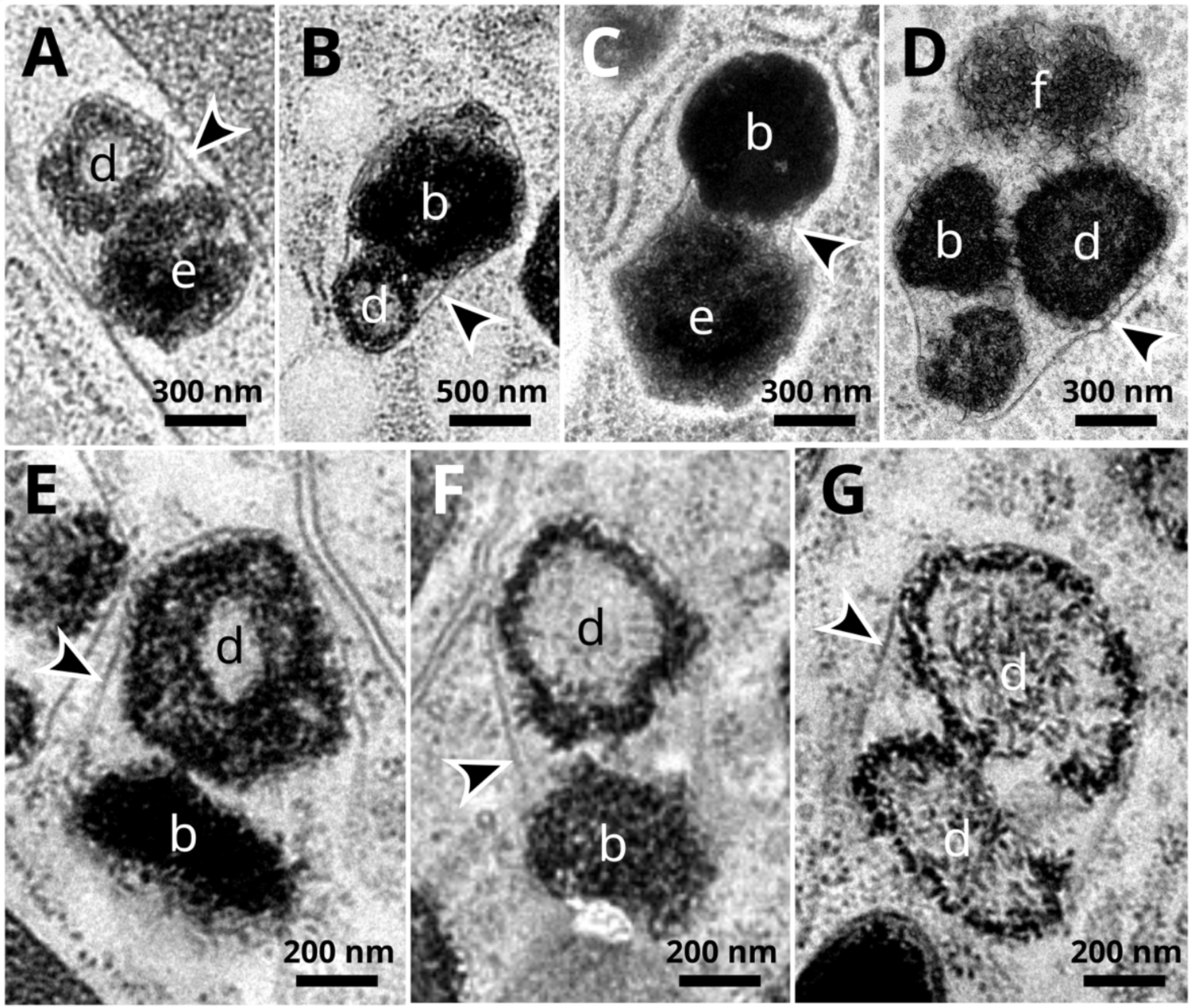
Various pigment organelle types observed within c-type clusters. (A-G) The various types of pigment organelles recognizable within c-type clusters are indicated by their respective letters. All c-types possess a single limiting membrane (arrowheads).

**Fig. S4.**
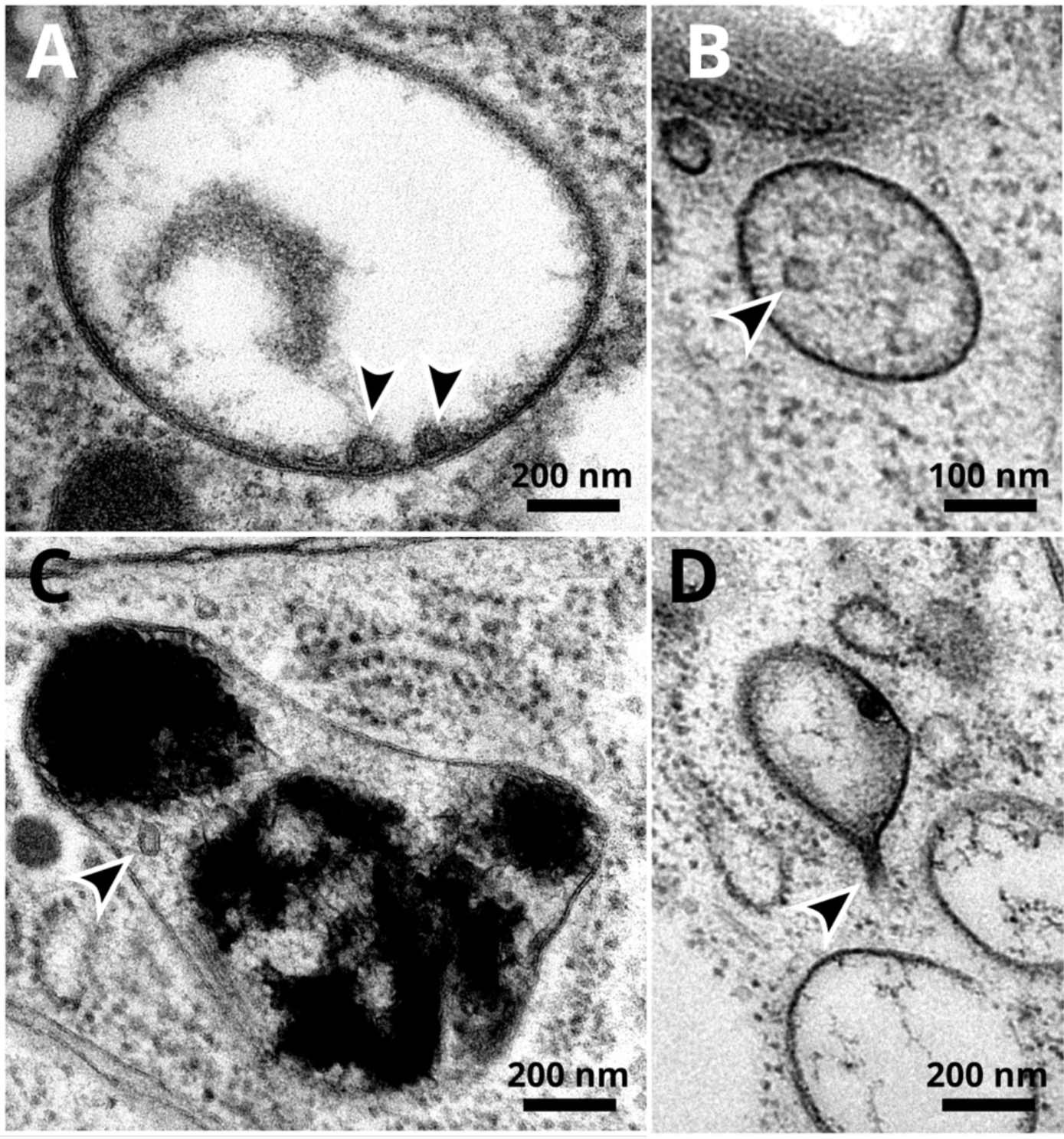
Intraluminal vesicles and membrane tubulations of pigment organelles. (A) Two intaluminal vesicles (arrowheads) closely associated to the limiting membrane of an early a-type pigment organelle. (B) Intraluminal vesicles and fibrils in a multivesicular body resembling a maturating a-type pigment organelle. (C) A intraluminal vesicle (arrowhead) within a c-type cluster of pigment organelles. (D) Tubulation of the limiting membrane (arrowhead pointing toward the tubule neck) of an early a-type pigment organelle.

**Fig. S5.**
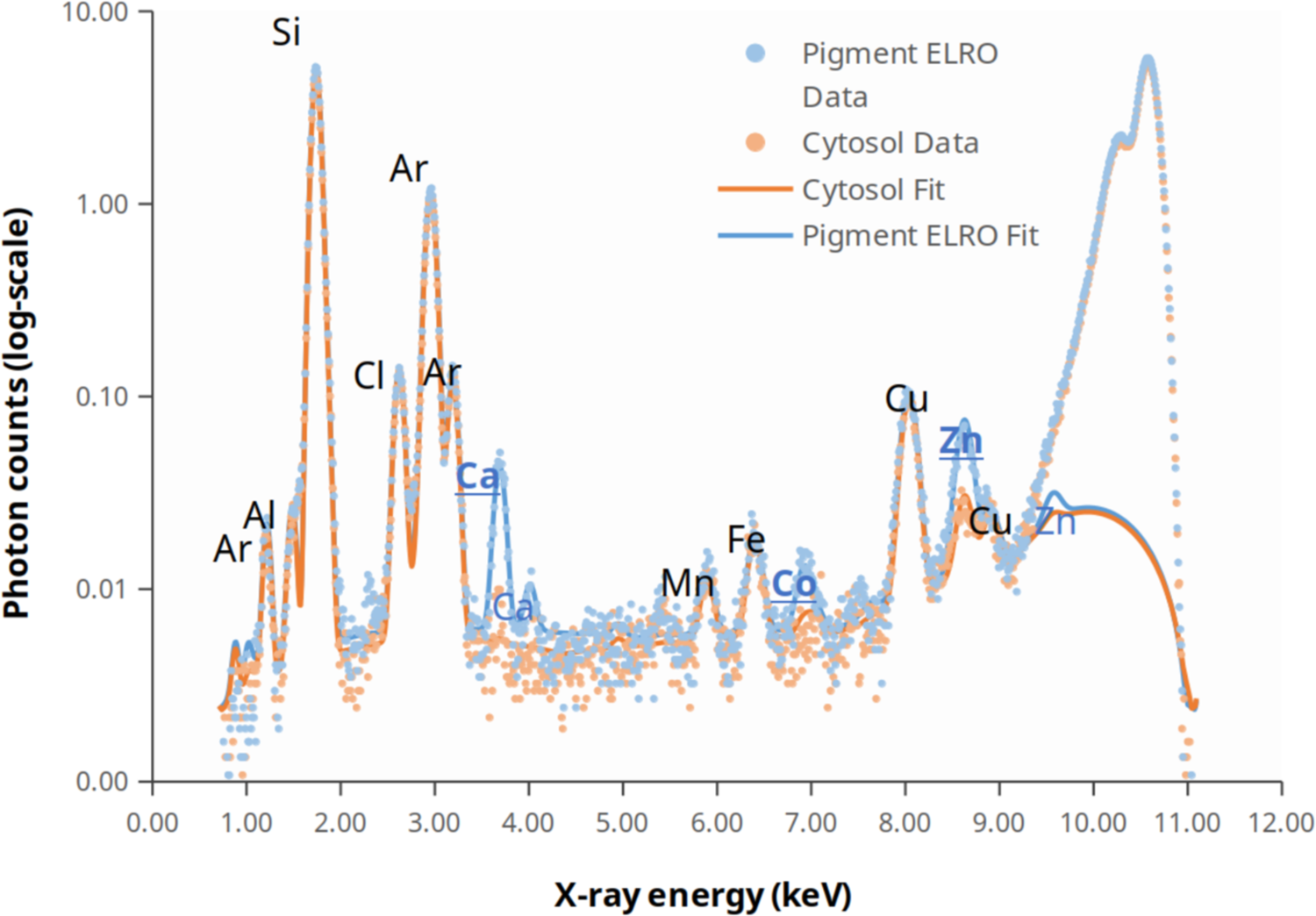
Recorded Synchrotron X-ray spectra of two intracellular regions within the same cell. X-ray fluorescence of native metals were obtained by exciting the area shown in the inset Figure 3C, which contains a single pigment organelle surrounded by cytosol. Recorded data are displayed as dots. Main fluorescence peaks (K-lines) of elements between Al and Zn are shown as continuous lines. X-ray energies above 9.7 keV are dominated by scattered X-rays. Elements indicated in blue fonts (Ca, Co and Zn) are those with higher fluorescence in the pigment organelle than in the cytosol. The peaks of underlined elements are those used for metal mappings in this study.

**Fig. S6.**
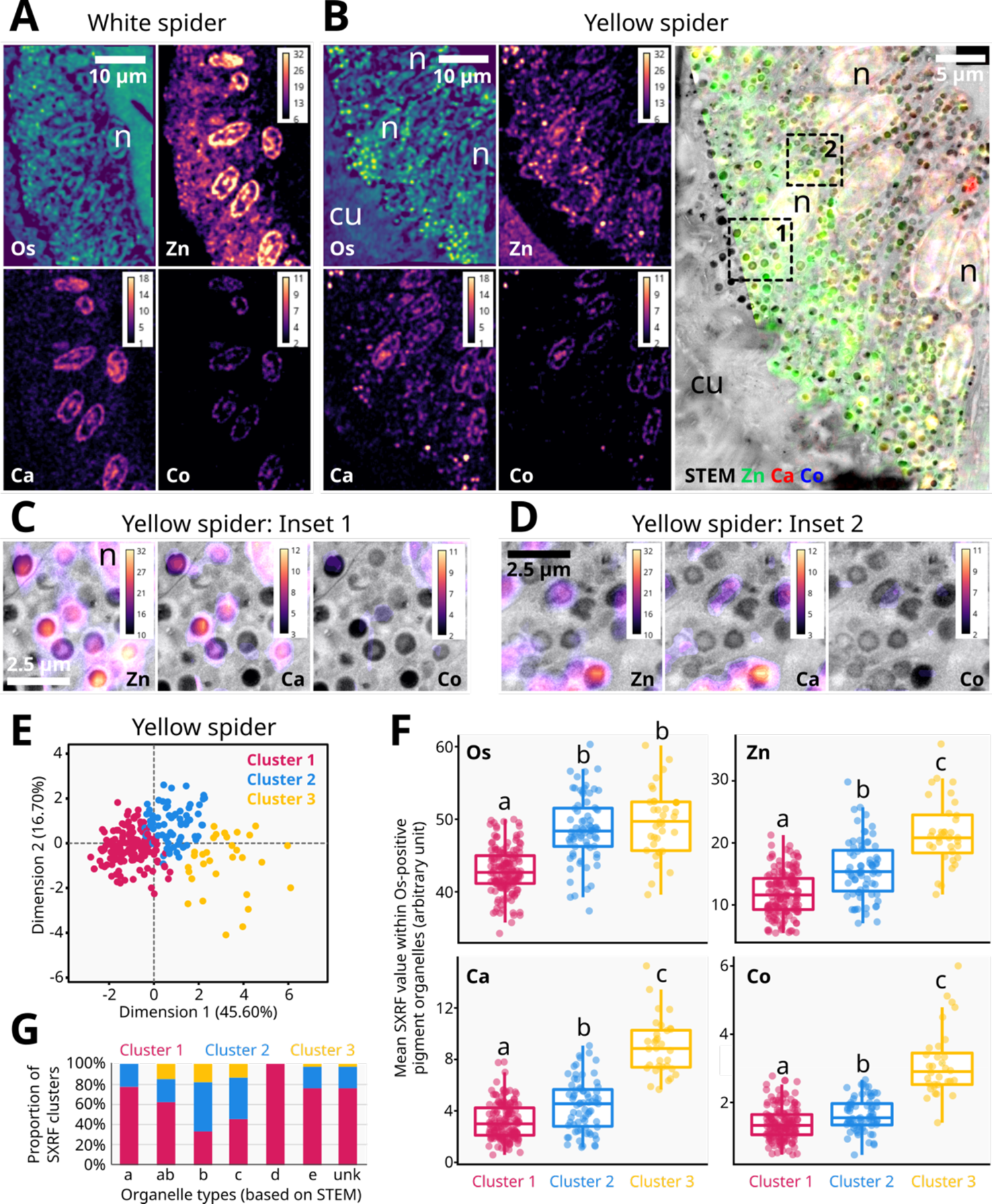
Pigment organelles differentially accumulate metals according to coloration and maturation stages. (A-B) Synchrotron X-ray fluorescence (SXRF) mapping of osmium (Os) and native zinc (Zn), calcium (Ca) and cobalt (Co) in white (A) and yellow (B) integuments. A correlative SXRF-STEM imaging of the yellow integument is provided. cu, cuticle; n, nucleus. (C-D) Zoom of the two regions depicted in B. (E) Principal component analysis and hierarchical clustering of Os-positive organelles in B based on their average content in Os, Zn, Ca, Co, S and Ni. (F) Metal contents in the clusters depicted in E. (G) Relative abundance of SXRF clusters defined in E according to organelle types. Different letters indicate statistical significance between clusters (p-value < 0.05; Kruskal-Wallis test followed by pairwise Wilcoxon rank sum test using Holm method for p-value adjustment). ab, pigment organelles with an intermediary morphology between a- and b-types; Unk, unknown organelles whose ultrastructure by STEM was too ambiguous to be classified.

**Fig. S7.**
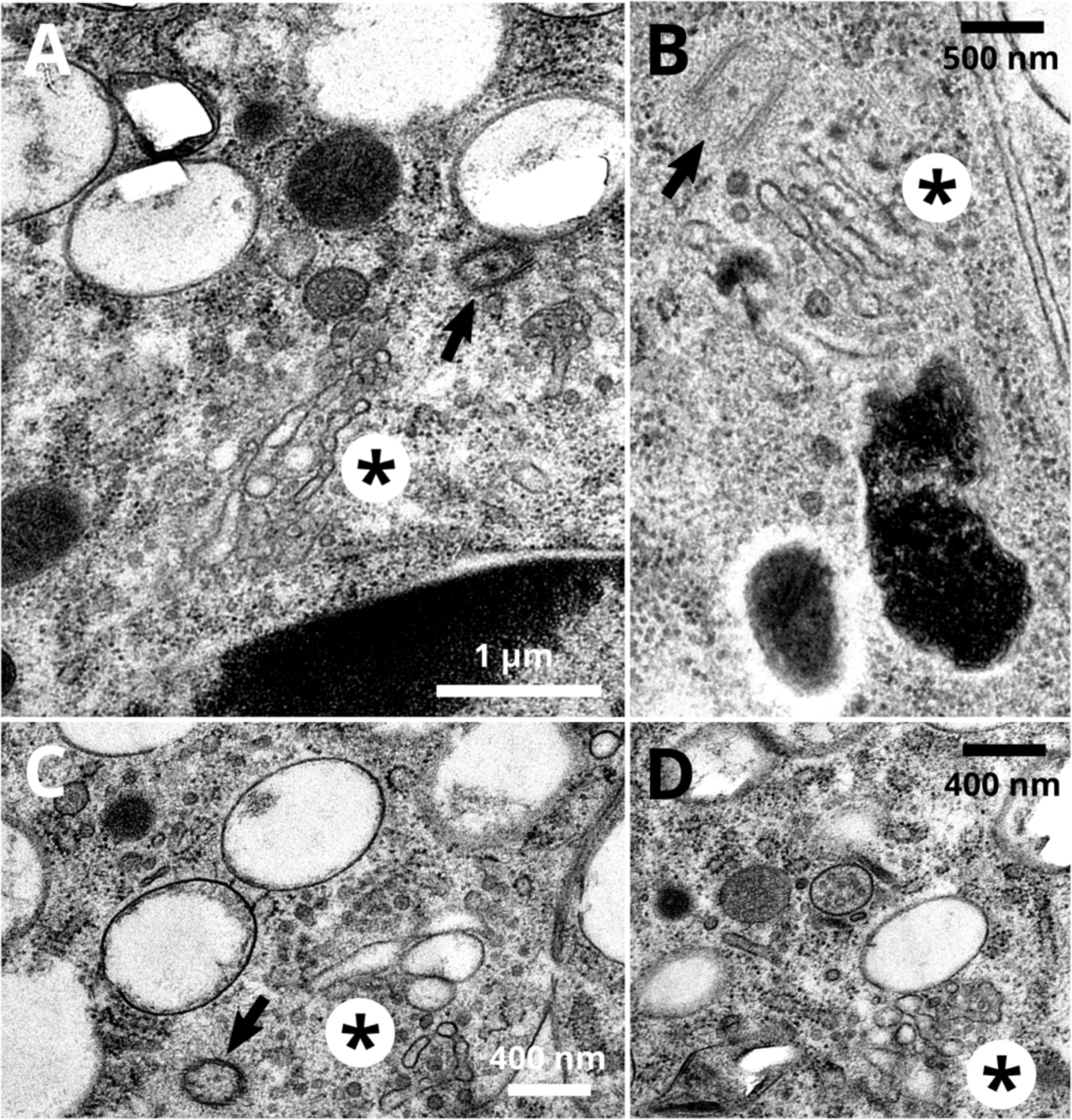
Examples of tubulo-saccular complexes in pigment cells of crab spiders. (A-C) Centrioles (arrows) can be observed in regions where tubulo-saccular complexes (asterisk) are situated. (D) Another example of a tubulo-saccular complex closely associated to an early a- type pigment organelle, endosomal compartments, vesicles and tubules.

**Fig. S8.**
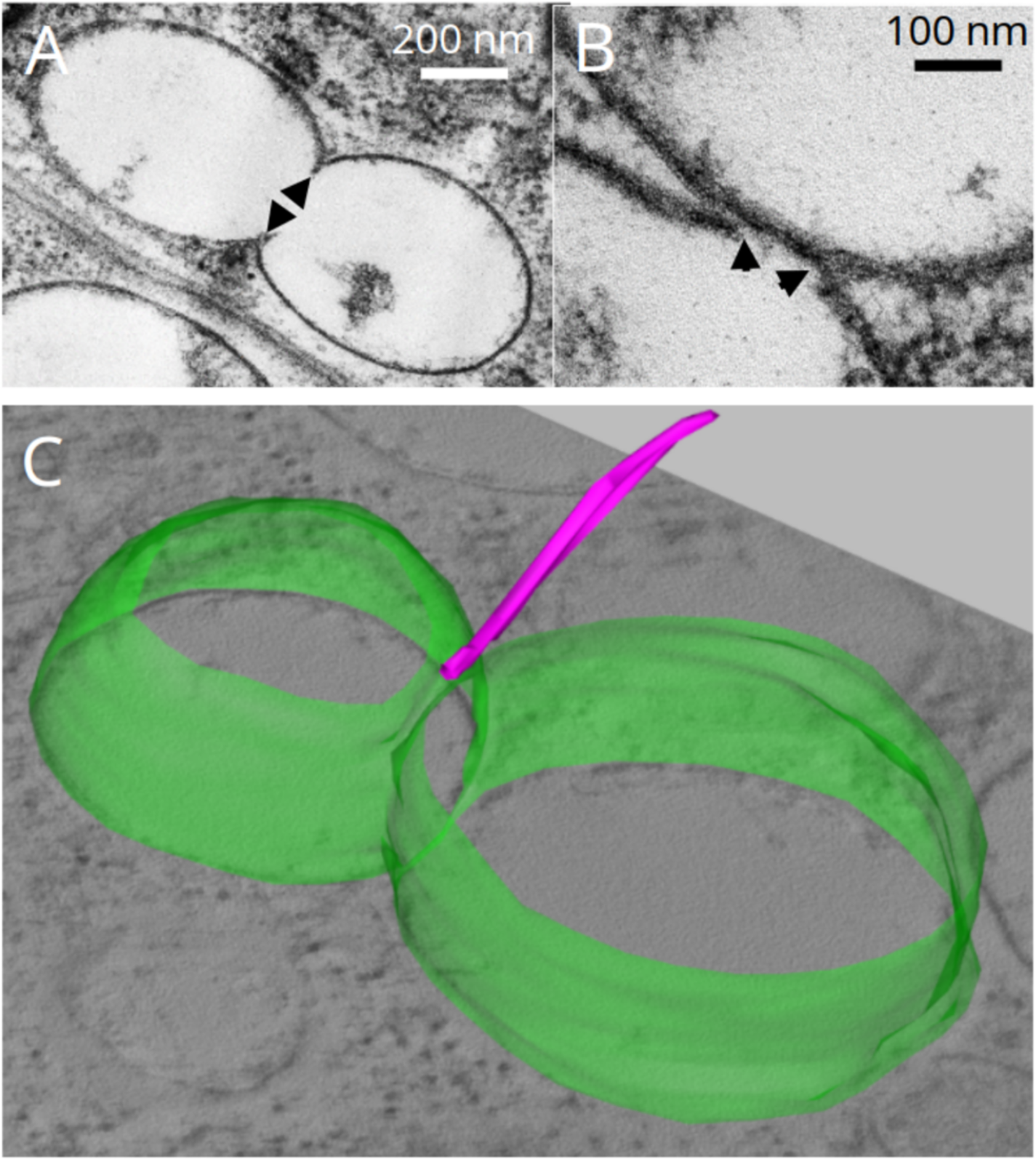
Evidence of homotypic fusion events between a-type pigment organelles in white spiders. (A-B) Continuity of limiting membranes (arrowheads) suggest homotypic fusion events between a-type pigment organelles. (C) Three-dimensional reconstruction of fused a-type pigment organelles (green membrane) showing a microtubule (magenta) closely-associated to the fusion site.

**Fig. S9.**
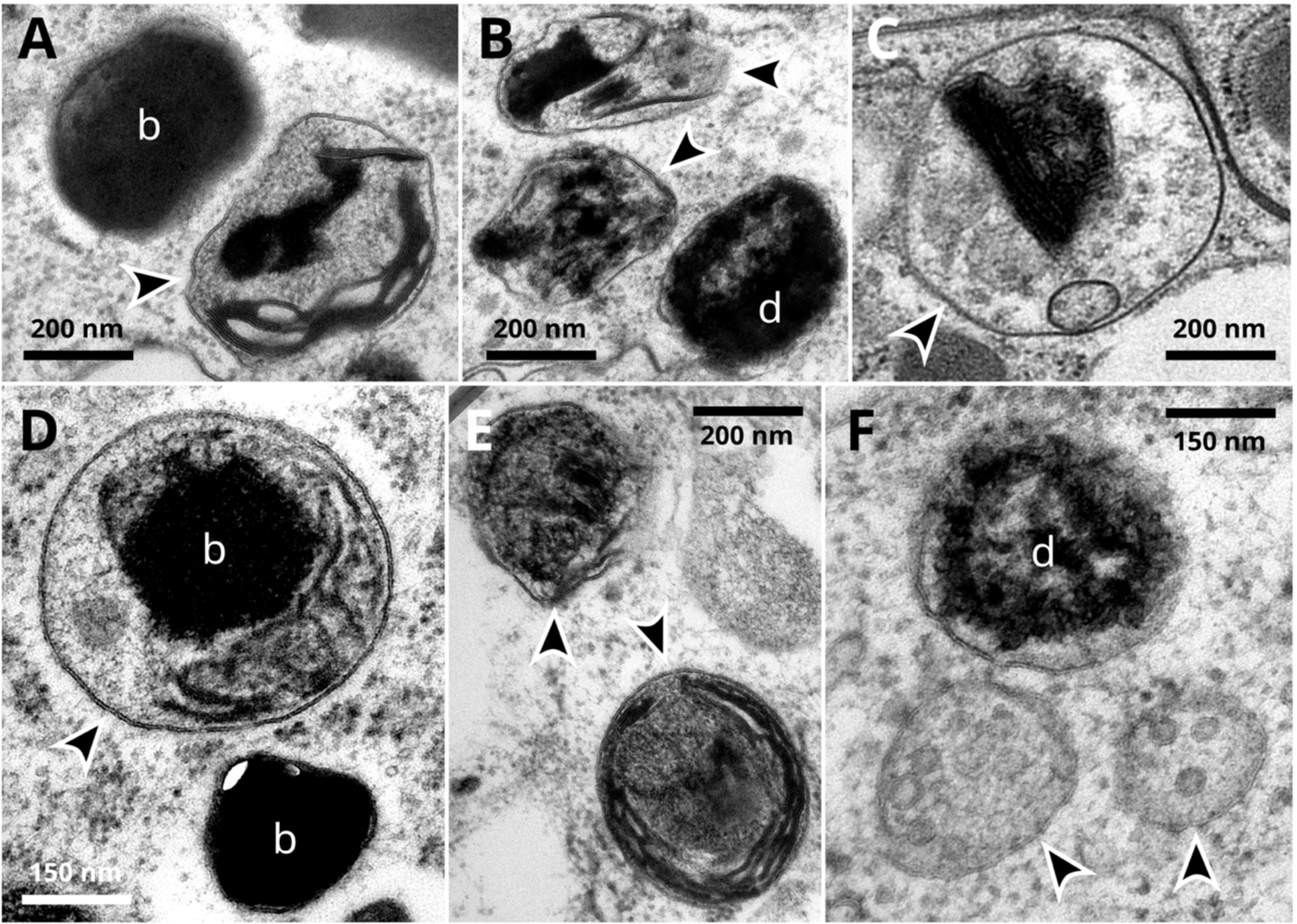
Ultrastructural evidence of lysosomal degradation of pigment organelles. (A-E) Lysosomal compartments (arrowheads) displaying a single limiting-membrane, dense lumens, heterogeneous and aggregated materials, as well as cores of b-type pigment organelles are observed, sometimes in the close vicinity of free pigment organelles. (F) Two multivesicular bodies (arrowheads) with dense granular lumens resembling late endosomes are closely associated to a d-type pigment organelle.

### Supplemental Tables

**Table S1.**
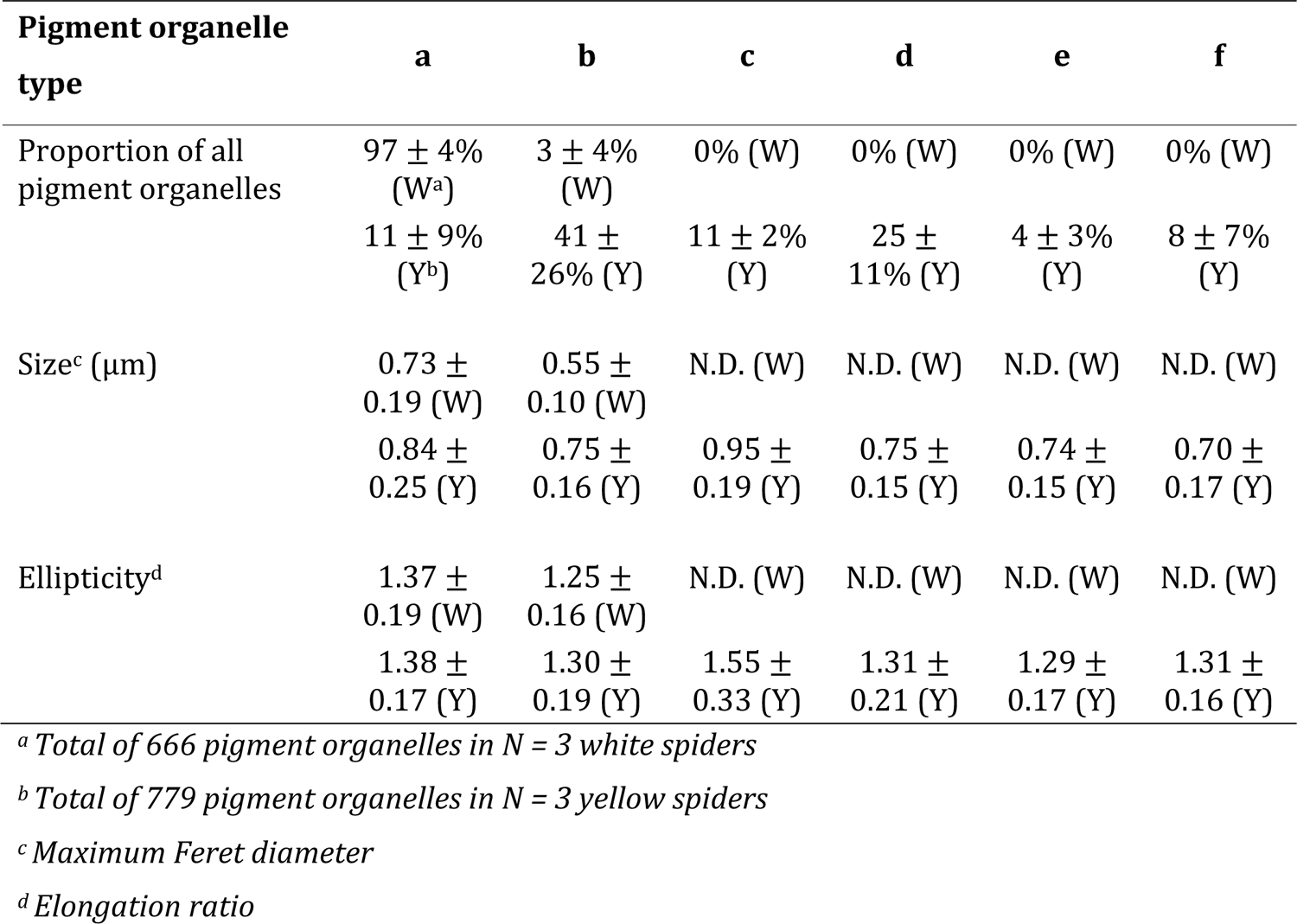
Characteristics of pigment organelle types in white (W) and yellow (Y) spiders (mean ± SD)

### Supplemental Movies

Movie S1. Tomographic reconstruction of a pigment organelle showing intraluminal vesicles (in blue) and sheet-like fibrils (in gold) associated to the limiting membrane (in green).

Movie S2. Tomographic reconstruction of a c-type pigment organelle showing several cores of intraluminal content (in gold) and two tubulations of the limiting membrane (in green).

Movie S3. Tomographic reconstruction of a tubulo-saccular complex (limiting membranes in various colors and intraluminal fibrillary content in gold) and its associated organelles, including free vesicles and tubules (in magenta), a-type pigment organelles (limiting membrane in green and intraluminal fibrillary content in gold) and an endosome (limiting membrane in purple and intraluminal vesicles in blue).

Movie S4. Tomographic reconstruction of the endosome (in purple) observed in Movie S3, displaying intraluminal vesicles (in blue) and a membrane tubulation.

Movie S5. Tomographic reconstruction of a d-type pigment organelle and its intraluminal content (in gold) displaying a pillar-like structure connecting the limiting membrane (in green).

Movie S6. Tomographic reconstruction of a f-type pigment organelle and its intraluminal content (in gold) showing its intertwined sheet-like structure.

Movie S7. Tomographic reconstruction of a c-type pigment organelle and its intraluminal content (in gold) displaying a pillar-like structure that connects its two cores.

